# The initiation and early development of apical-basal polarity in *Toxoplasma gondii*

**DOI:** 10.1101/2024.07.14.603470

**Authors:** Luisa F. Arias Padilla, Jonathan Munera Lopez, Aika Shibata, John M. Murray, Ke Hu

## Abstract

The human parasite *Toxoplasma gondii* has a distinctive body plan with a well-defined polarity. In the apical complex, the minus ends of the 22 cortical microtubules are anchored to the apical polar ring, a putative microtubule-organizing center. The basal complex caps and constricts the parasite posterior end, and is critical for cytokinesis. How this apical-basal polarity axis is initiated was unknown. Here we examined the development of the apical polar ring and the basal complex in nascent daughters using expansion microscopy. We found that different substructures in the apical polar ring have different sensitivity to stress. In addition, apical-basal differentiation is already established upon nucleation of the cortical microtubule array: arc forms of the apical polar ring and basal complex associate with opposite ends of the microtubules. As the construction of the daughter framework progresses towards the centrioles, the apical and the basal arcs co-develop in striking synchrony ahead of the microtubule array, and close into a ring-form before all the microtubules are nucleated. We also found that two apical polar ring components, APR2 and KinesinA, act synergistically. The removal of each protein individually has modest to no impact on the lytic cycle. However, the loss of both results in abnormalities in the microtubule array and highly reduced plaquing and invasion efficiency.

## INTRODUCTION

The ∼ 6000 known members of the phylum Apicomplexa are unicellular parasites that infect a wide range of invertebrate and vertebrate hosts [1]. Many of them cause devastating diseases in humans, such as cryptosporidiosis, toxoplasmosis, and malaria [1–4]. The apicomplexans are also excellent models for exploring the construction, function, and evolution of cellular architectures, due to their sophisticated membrane and cytoskeletal structures and *de novo* generation of a membrane-cytoskeletal scaffold for daughter cells during cell division [5–11] (Figure 1).

**Figure 1.**
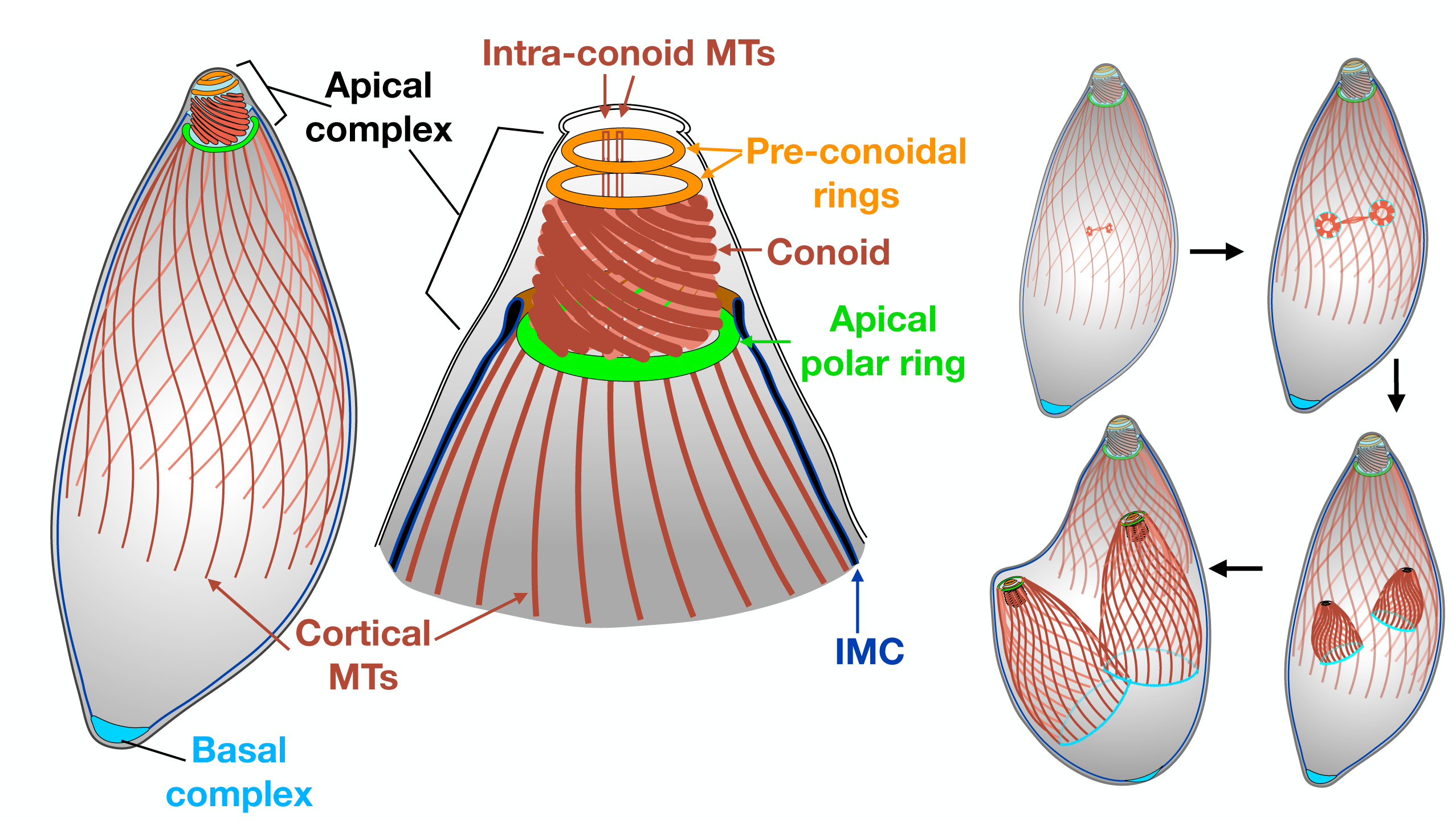
Schematic of tubulin-containing and associated structures (*left*) and daughter formation throughout the cell cycle (*right*). For clarity, the cortical microtubules (MTs) of the mother are shown in a lighter color. Membrane-bound organelles are not shown. IMC: Inner membrane complex.

In *Toxoplasma gondii,* the two poles of the parasite are marked by the apical and basal complexes, respectively [12–14] (Figure 1). The cytoskeletal elements of the apical complex include the apical polar ring, the conoid and associated preconoidal rings and the two intra-conoid microtubules (MTs) [12, 15, 16]. The conoid is formed of highly curved, ribbon-like tubulin fibers [16–19]. The entire conoid protrudes or retracts through the apical polar ring in a calcium-dependent manner. The apical polar ring is also structurally integrated with the minus ends of the 22 cortical MTs in mature parasites [16–18, 20, 21]. The removal of two apical polar ring components (KinesinA and APR1) results in MT detachment from the apex in mature parasites [22]. However, it is unknown how this putative MT-organizing center is assembled and how it develops with respect to the MT array.

The basal complex caps and constricts the posterior end of the parasite [13, 14, 23–25]. Although the sophistication of the apical complex is immediately obvious from its striking structure and has been known for many decades, the complexity of the basal complex was not recognized until much later when multiple components of the basal complex were identified in a proteomic screen [13]. That screen was designed to identify components of the apical complex, but unexpectedly a number of the apical complex components were also localized to the basal complex. One of them, MORN1, is weakly localized to the apical complex, but has a very prominent localization to the basal complex and the spindle pole. Later, MORN1 was found to be important for maintaining the constriction of the parasite posterior end and successful cytokinesis [23, 25]. The basal complex was first discovered as a ring abutting the distal end of the cortical MTs in daughter parasites [13, 14]. After the daughters emerge from the mother parasite, the basal complex disengages from the cortical MTs via an unknown mechanism and becomes the basal cap of the mature parasite. Although MORN1 and some other components of the basal complex have been shown to be recruited to early daughters [14, 26], how the basal complex initiates and develops into a ring was not known due to the limited temporal and spatial resolution of existing microscopy data.

Recently, we investigated the initiation and early development of the tubulin-containing structures in *Toxoplasma* using expansion microscopy [27]. We discovered that the 22 cortical MTs are not initiated at the same time. The daughter MT arrays first are detected close to the duplicated centrioles as incomplete cogwheels, in which 6 to 7 “stubs” of MTs form around the primordial conoid. The assembly of the MT array proceeds towards the centrioles. With the elongation and addition of more MTs, “petals” that contain 4 or 6 MTs form in the disc-like nascent daughters. After all 22 MTs are nucleated, the 5-fold symmetry of the early MT array becomes established, often with a 4, 4, 4, 4, and 6 grouping pattern. This study uncovered distinct structural intermediates during nucleation, elongation, and symmetry development of the array. It also raised the question as to how the cytoskeletal complexes that associate with the opposite ends of the MT array develop, *i.e.*, do they follow a concerted or disjoint assembly path with respect to each other and to the MT array?

Here we address these questions by delineating the localization of components of the apical polar ring (APR2) and the basal complex (MORN1) using expansion microscopy. The results reveal the extremely early establishment of the apical-basal polarity and a striking co-development of the arc-form precursors of the apical polar ring and the basal complex at opposite ends of the cortical MTs. We also found that different structural domains in the apical polar ring have different sensitivity to stress. Lastly, we examined the impact of loss of APR2 alone and with KinesinA, and found that these two proteins act synergistically in plaquing efficiency, MT organization, and parasite invasion.

## RESULTS

### The apical polar ring develops ahead of the cortical MT array during the initial assembly of the array

Previously, we carried out a comparative proteomic screen to identify protein components of the apical complex of *Toxoplasma* [13]. For one of the proteins, TgGT1_227000, 10 peptides and 16 spectra were captured in the apical complex enriched-fraction and no peptides were found in the apical complex-depleted fraction. Confirming the highly significant enrichment in the apical complex-fraction in our work, TgGT1_227000 was also identified as one of the apical complex proteins in a more recent BioID screen [28]. In that study, TgGT1_227000 was localized to the apical polar ring of the mature parasites, but its localization in the daughter parasites was not determined.

To investigate the localization of this protein during daughter development, we generated a knock-in parasite line in which TgGT1_227000 (now named as the **a**pical **p**olar **r**ing protein **2** or **APR2**) is endogenously tagged with mEmerald (Figure 2A). Three-dimensional structured illumination microscopy (3D-SIM) confirmed the localization to the apical polar ring in mature parasites (Figure 2B, arrowheads). Additionally, APR2 is detected at the apex of nascent daughters, before the dome-like shape of the daughters can be discerned (Figure 2B, white arrows). This provides an opportunity to answer critical questions related to the spatial and temporal relationship between the development of the apical polar ring and the cortical MT array: Does the apical polar ring assemble into a ring prior to nucleation of the MTs or does it co-develop with the MT array?

**Figure 2.**
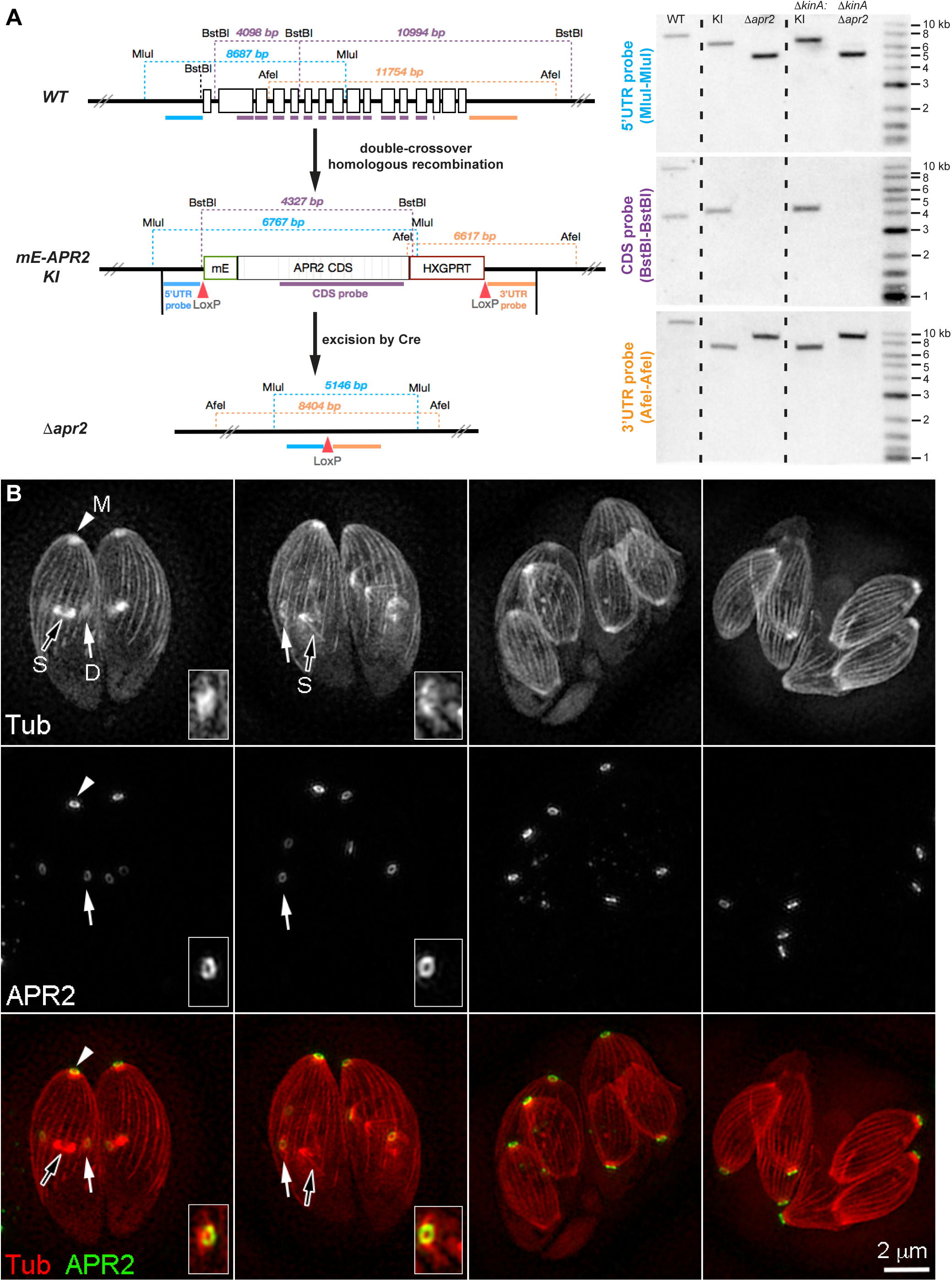
Generation of *mEmeraldFP-APR2* knock-in and *Δapr2* parasites and localization of APR2 in live parasites. **A**. *Left:* Schematic for generating the *mEmeraldFP-APR2* knock-in (*mE-APR2* KI), *Δapr2* parasites and Southern blotting strategy. Restriction sites, hybridization sections of the Southern blot probes for the *apr2* coding region (CDS, purple bar), regions upstream (“5’ UTR probe”, blue bar) and downstream (“3’ UTR probe”, orange bar) of CDS, and the corresponding DNA fragment sizes expected are indicated. Vertical black bars in the *mE-APR2* KI schematic indicate the boundary of the sequences of 5’ and 3’UTR included in the *pTKO2_II-mE-APR2* knock-in plasmid used for generating the *mE-APR2* KI parasite. *Right:* Southern blots of the RH*Δku80* parental (WT), *mE-APR2* knockin (KI), *Δapr2*, *ΔkinesinA*: *mE-APR2* knockin (*ΔkinA*:KI), and *ΔkinesinAΔapr2* (*ΔkinAΔapr2*) parasites. Τhe hybridization patterns confirmed the homologous integration of the mE-APR2 fusion in the KI and *ΔkinesinA*:KI parasites, and the deletion of the *apr2* locus in the *Δapr2* and *ΔkinesinAΔapr2* parasites. The expected patterns are described in Materials and Methods. **B.** Projections of 3D-SIM images of live *mE-APR2* knock-in *Toxoplasma* parasites expressing mAppleFP-β tubulin 1, in which cortical MTs are fluorescently labeled. mE-APR2 is detected in nascent daughters forming close to the centrioles (first and second columns). White arrows and insets (2X): daughter (D) structures. Arrowheads: mother apical complex (M). Black arrows: spindle (S).

The resolution of 3D-SIM imaging is insufficient to resolve the detailed structures of the apical polar ring and the MT array. We therefore examined the assembly of the apical polar ring and the MT array using expansion microscopy (Figure 3). The *mEmeraldFP-APR2* (*mE-APR2*) knock-in parasite was labeled with an anti-GFP and an anti-tubulin antibody. We found that the assembly of the apical polar ring is initiated prior to that of the MT array. The precursor of the ring is first detected as an arc in the vicinity of the centrioles before the MT array is detectable (Figure 3A). Similar to the MT array and the conoid, the growth of the apical polar ring also progresses towards the centrioles. Interestingly, the arc of the APR2 labeling often extends beyond the forming MT array towards the centrioles (arrowheads, Figure 3 A-G). The apical polar ring has taken on its final ring-like shape before the completion of the nucleation of all 22 cortical MTs and before full establishment of the MT “petals” (Figure 3G-I). This, for the first time, reveals how the growth of the apical polar ring might direct that of the MT array. It is conceivable that the leading extension of the apical polar ring guides the initiation of new MTs by laying down or recruiting new nucleation complexes. Together, the development of both the apical polar ring and the MT array reveals a clear temporal gradient. MTs closer to the centriole region are younger; i.e., constructed later.

**Figure 3.**
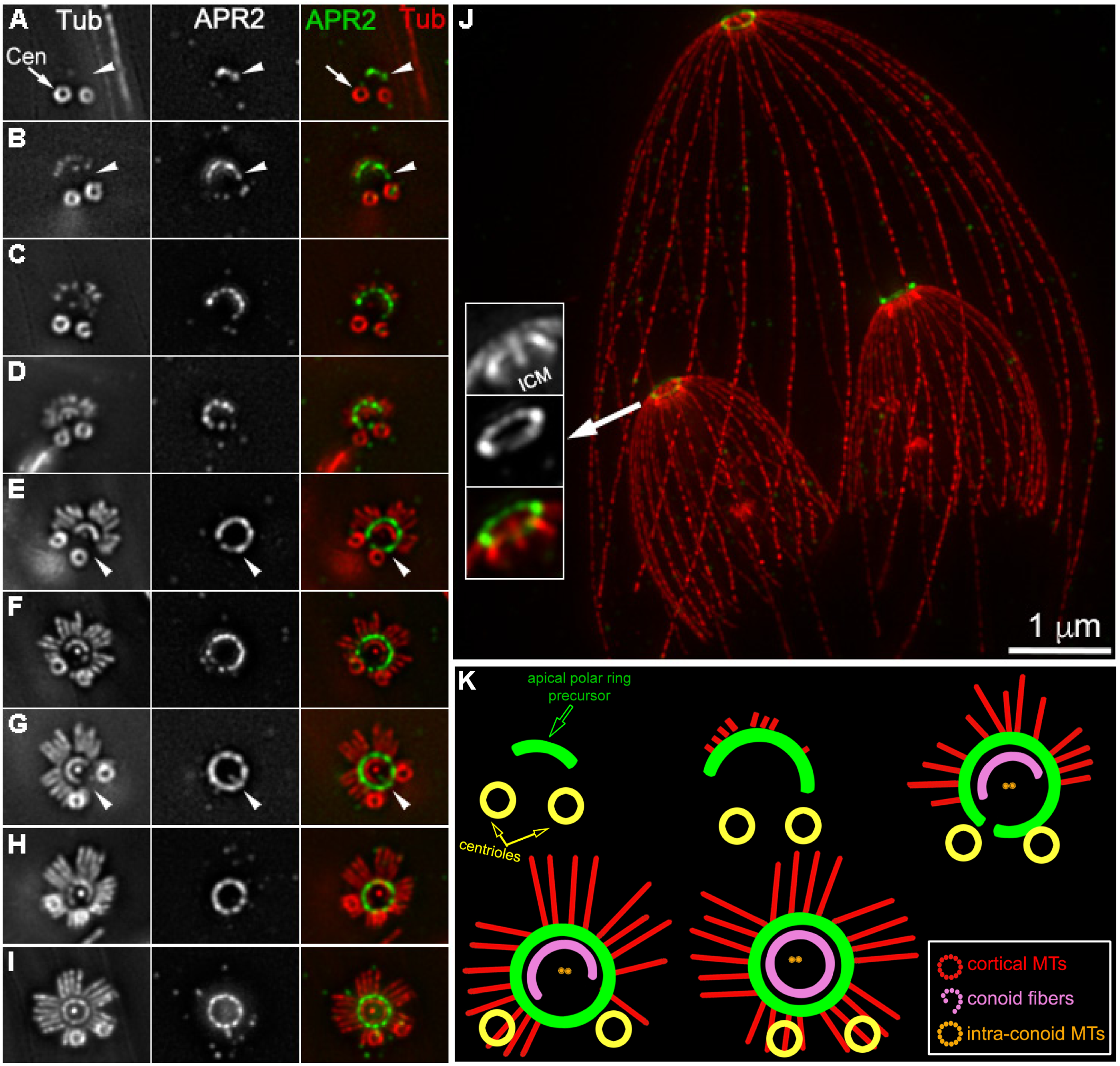
The apical polar ring extends ahead of the cortical MT array during daughter development **A-J.** Projections of expansion microscopy (ExM) images of *mE-APR2* knock-in *Toxoplasma* parasites labeled with anti-tubulin (grayscale and red) and anti-GFP (grayscale and green) antibodies. Insets (2X) in J are projections of a subset of the sections showing that the apical polar ring labeling is positioned apically to the retracted daughter conoid. ICM: Intra-<u>c</u>onoid <u>M</u>Ts. Image contrast was adjusted to optimize display. Arrowheads indicate sections of the apical polar ring that extend towards the centriole region beyond the forming MT array. Cen: centrioles. Bar ≈ 1 µm prior to expansion. **K.** Diagrams illustrating the development of cortical MTs (red), conoid (pink), intra-conoid MTs (orange) with respect to that of the apical polar ring (green) in the vicinity of the centrioles (yellow). Diagrams were drawn based on images in panels A, B, E, G and I.

### The basal complex develops in concert with the apical polar ring

We discovered previously that markers for both the apical structures and the basal complex are recruited to early daughters [13, 14]. The earliest discernible basal complex structure takes the form of a ring, which first expands, then constricts, and eventually forms a basal capping structure in the mature parasite [14]. However, how the apical-basal axis in nascent daughters is established during MT development was not known. Also unknown was the path that the basal complex follows to form into a ring. To address this, we examined the assembly of the basal complex with respect to that of the apical polar ring and the tubulin-containing structures using expansion microscopy. We and others previously discovered that MORN1 is one of the major components of the basal complex [13, 14, 23–25]. Using a MORN1 antibody, we labeled the basal complex at various stages of daughter formation (Figure 4). We found that the apical complex (including the conoid and the apical polar ring), the cortical MTs, and the basal complex develop in concert. In the earliest detectable daughters, where only 6-7 stubs of newly nucleated cortical MTs are present, the primordial basal complex is assembled as an arc at the outer circumference of the MT array, while at the same time the near concentric arc of the primordial apical polar ring is assembled at the inner circumference of the array (Figure 4A). As additional MTs are nucleated, both the apical and basal arcs extend towards the centrioles, just like the MT array and the conoid. In the meantime, the distance between the two arcs increases as the cortical MTs grow in length. While the radius of the apical polar arc/ring remains nearly constant throughout the assembly process, that of the basal complex steadily increases as the MTs grow outwards. The open form of the basal complex in early daughters was also recently reported in [29] when this manuscript was in preparation. Similar to the apical polar ring, the extension of the basal complex leads that of the MT array (Figure 4A-E, arrowheads). The closure of the MORN1 ring occurs before all 22 MTs are nucleated (Figure 4E). This suggests that the development of the basal complex is likely independent of that of the MT array, even though the assembly of these two sets of structures is highly coordinated. Of note, when ectopically expressed in bacteria, MORN1 can form ring-like structures, indicating that it has an intrinsic ring-forming propensity [23]. It is therefore conceivable that the basal ring formation is at least partly self-driven by core components such as MORN1, but the localization, timing and direction of assembly might be modulated by proteins that couple the basal complex to the MT array.

**Figure 4.**
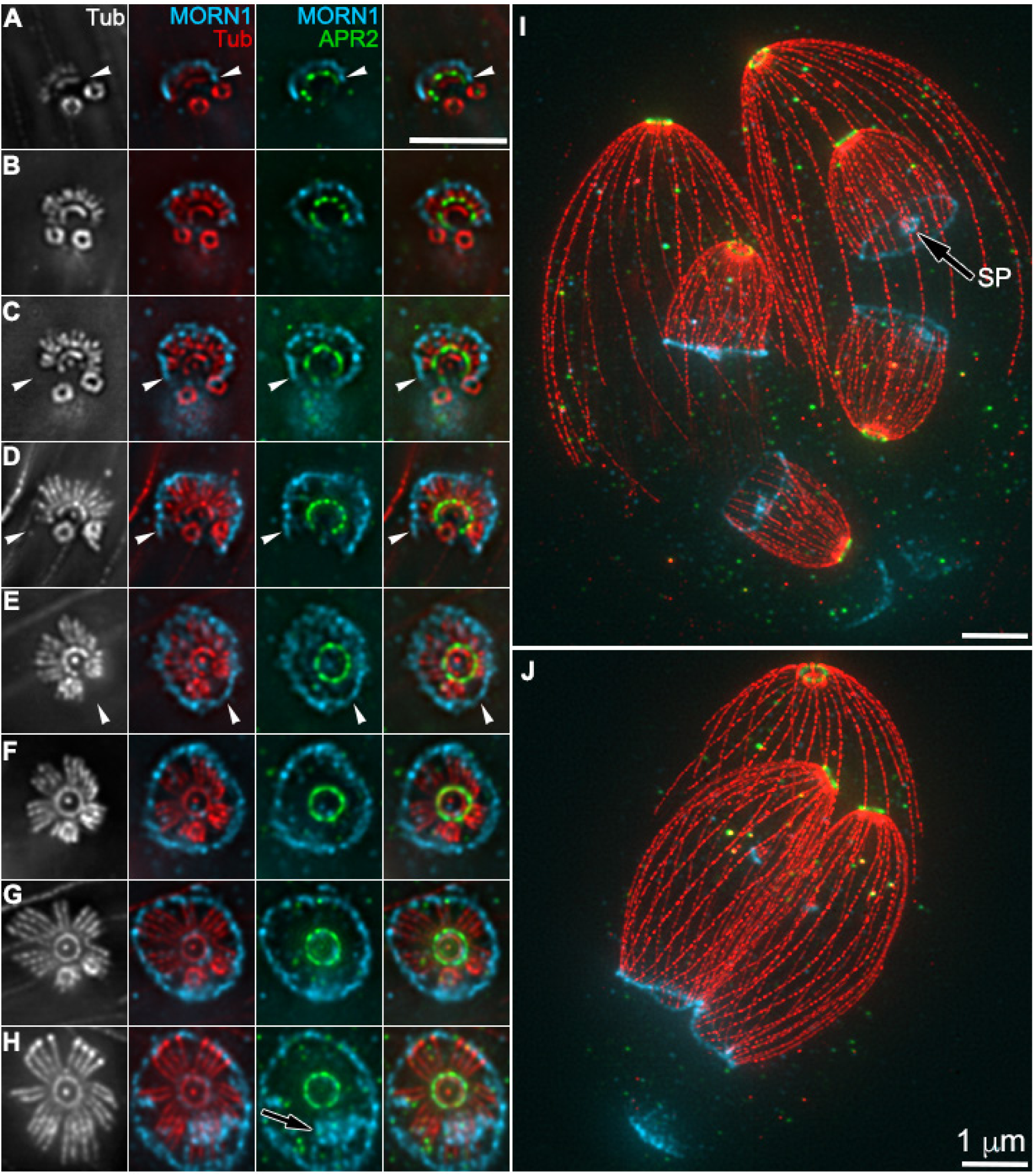
The basal complex co-develops with the cortical MT array and the apical polar ring. **A-J.** Projections of ExM images of *mE-APR2* knock-in *Toxoplasma* parasites labeled with anti-tubulin (grayscale and red), anti-GFP (green), and anti-MORN1 (cyan) antibodies. Arrowheads indicate sections of the basal complex that extend towards the centriole region beyond the forming MT array. Black arrows: spindle pole (SP). Image contrast was adjusted to optimize display. Bars ≈ 1 µm prior to expansion.

Previously we generated a triple knockout mutant (denoted “TKO”) in which three MT associated proteins, TLAP2, TLAP3, and SPM1 were removed [30]. The cortical MTs are destabilized in mature TKO parasites. The destabilization is particularly exacerbated in dividing parasites [31]. Here we found that the development of the daughter apical polar ring, and the basal complex are not affected in the TKO parasite (Figure 5A-B). Often only a collar of tubulin surrounding the apical polar ring is left at the apex of the mother parasite (Figure 5B, insets), which results in a striking, exposed view of the daughter frameworks in the full projection of the parasite.

**Figure 5.**
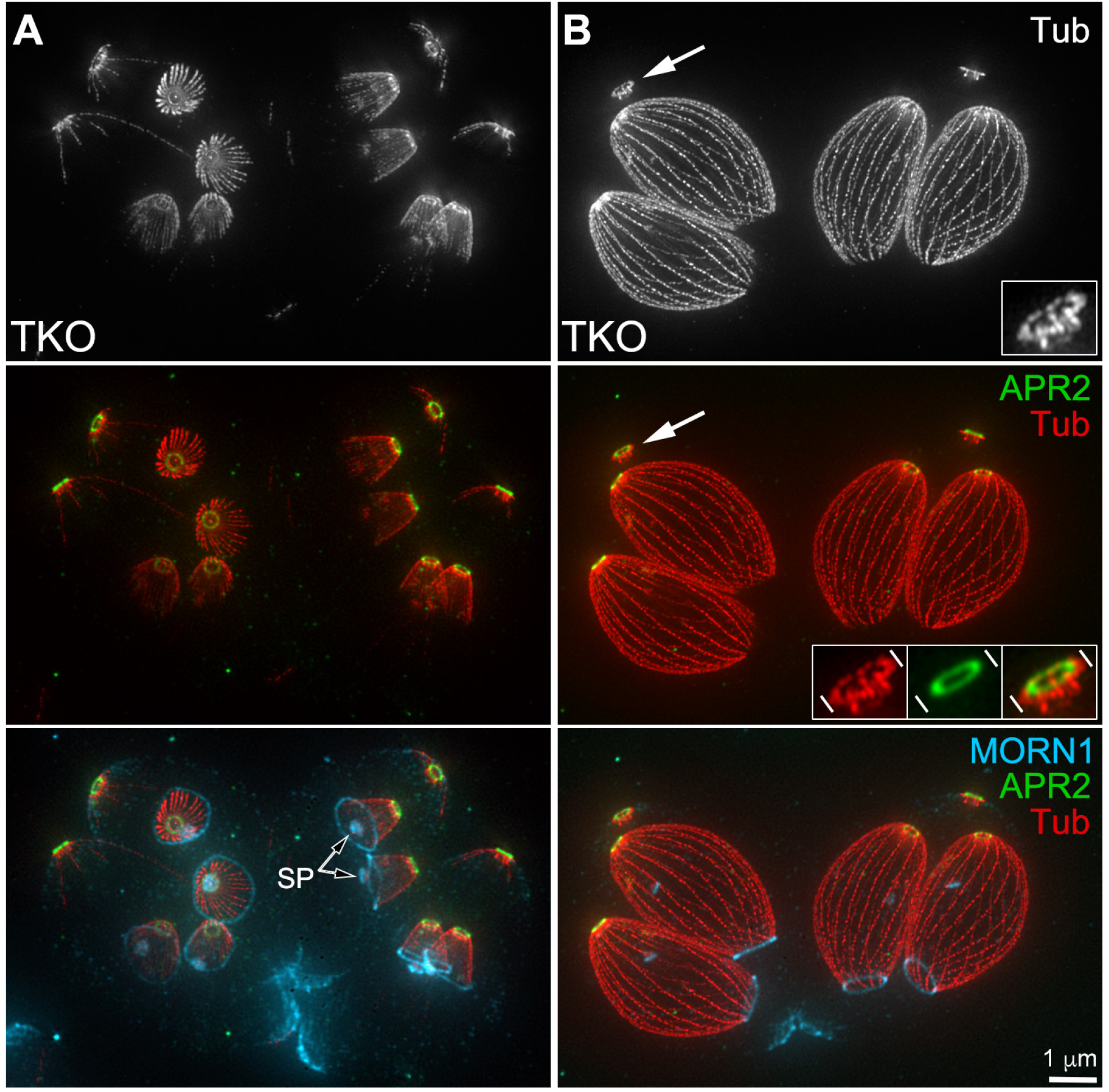
The basal complex, cortical MT array and the apical polar ring assemble normally in the TKO parasites. Projections of ExM images of TKO parasites expressing mE-APR2 labeled with anti-tubulin (grayscale and red), anti-GFP (green), and anti-MORN1 (cyan) antibodies. Black arrows: spindle pole (SP). The panels include vacuoles containing parasites at an earlier (A) and a late stage (B), respectively. Insets: 3X enlargement of the parasite apex indicated by the white arrows, in which a remnant tubulin “collar” around the apical polar ring is seen. Image contrast was adjusted to optimize display. Bar ≈ 1 µm prior to expansion.

### The apical polar ring has at least two molecularly distinct substructures with different sensitivity to stress

Previously we identified another early component of the apical polar ring, KinesinA. To determine the interdependency of early components of the apical polar ring, we generated a *ΔkinesinA*: *mE-APR2* knock-in line, and found that APR2 remains associated with the apical polar ring in the absence of KinesinA (Figure 6A). Therefore APR2 localization is independent of that of KinesinA. When KinesinA was knocked out together with APR1, a later component of the apical polar ring, the cortical MTs detach from the apex in mature parasites [22]. The labeling of RNG1, another apical polar ring component [32], also fragments in the *ΔkinesinAΔapr1* parasites. To determine if APR2 localization is affected the same way, we generated a *ΔkinesinAΔapr1*: *mE-APR2* knock-in line, and found that mE-APR2 labeling remains as a ring in the *ΔkinesinAΔapr1* parasite even when the MT dispersal from the apex is evident (Figure 6B). To directly compare the RNG1 and APR2 localization, we generated RNG1-mCherry expressing lines in both the WT and *ΔkinesinAΔapr1*: *mE-APR2* knock-in background. We found that in the wild-type parasite, APR2 and RNG1 labelings are distinct, with the RNG1 ring being wider and more basal. In the *ΔkinesinAΔapr1* parasite, while the labeling of RNG1 fragments, APR2 labeling still retains the ring-shape in the same parasite (Figure 6C). This indicates that APR2 and RNG1 belong to two distinct structural domains that have different sensitivity to stress imposed on the apical polar ring during parasite growth.

**Figure 6.**
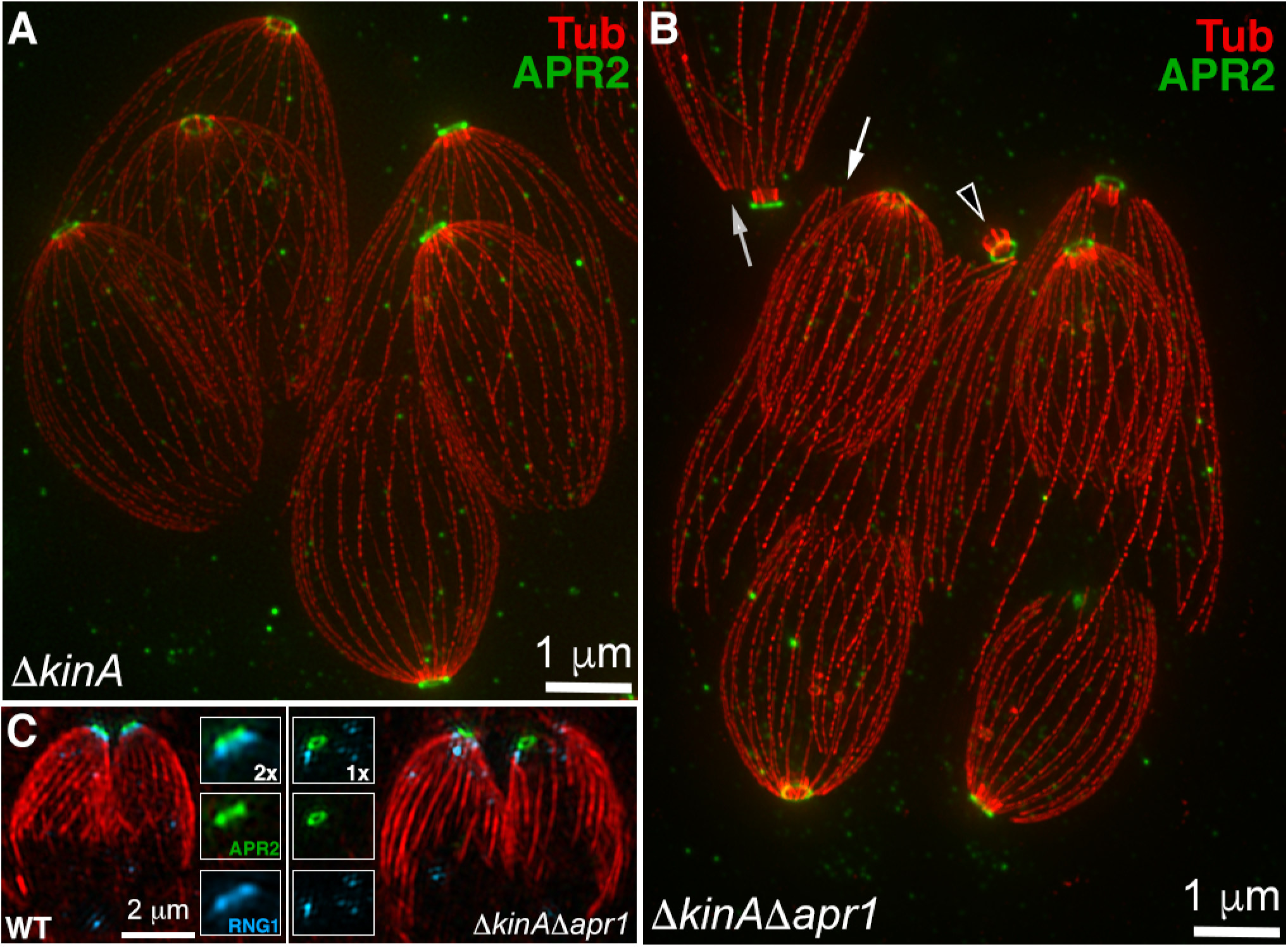
The apical polar ring has at least two molecularly distinct structural domains. **A.** Projections of ExM images of *ΔkinesinA*: *mE-APR2* knock-in parasites labeled with anti-tubulin (red) and anti-GFP (green) antibodies. Bar ≈ 1 µm prior to expansion. **B.** Projections of ExM images of *ΔkinesinAΔapr1*: *mE-APR2* knock-in parasites labeled with anti-tubulin (red) and anti-GFP (green) antibodies. White arrows: detached MTs. Arrowhead: detached conoid. **C.** Projections of SIM images of *mE-APR2* knockin and *ΔkinesinAΔapr1*: *mE-APR2* knockin parasites expressing RNG1-mCherry. Red: Anti-tubulin. Green: mE-APR2. Cyan: RNG1-mCherry.

### Loss of APR2 alone does not have a detectable impact on the patterning of the cortical MTs or parasite lytic cycle, but the loss of both APR2 and KinesinA has a synergistic effect

To determine the structural impact of APR2 on the apical polar ring and MT organization, we generated an APR2 knockout line - *Δapr2,* by transiently expressing Cre recombinase in the *mE-APR2* knockin parasite to excise the LoxP-mE*-*APR2-[selectable marker] cassette, generating individual clones that had lost the mE-APR2 fluorescence, and then confirming the excision of the APR2 locus by Southern blots using probes specific to the APR2 CDS as well as noncoding regions upstream (5’) and downstream (3’) from the CDS (Figure 2A). To determine how combined loss of APR2 and KinesinA affects the patterning and development of the cortical MT array, we also generated *ΔkinesinAΔapr2* parasites from the *ΔkinesinA*: *mE-APR2* knockin line by Cre-LoxP based excision as described for *Δapr2* (Figure 2A).

We examined the number and arrangement of the MTs in the array in the wild-type parental line (RH*ΔhxΔku80-*denoted “RH*Δku80*” or WT in what follows), *ΔkinesinA*, *Δapr2*, and *ΔkinesinAΔapr2* parasites using expansion microscopy and anti-tubulin immunofluorescence (Figure 7A-B). MT detachment or severe MT disorganization was detected in 0% of the mature WT parasites (total 727 parasites counted in three independent experiments), 2% of the *ΔkinesinA* (722 counted), 0.4% of the *Δapr2* (545 counted), and 10% of the *ΔkinesinAΔapr2* (567 counted) parasites. The impact of the double knockout of KinesinA and APR2 is therefore more pronounced than the additive effect of the single knockouts, but it is still quite mild. Much more severe MT detachment occurs in the *ΔkinesinAΔapr1* parasite (86.6%, 156 counted), indicating that APR1 plays a more prominent role in reinforcement of the apical polar ring than APR2 (Figure 7A).

**Figure 7.**
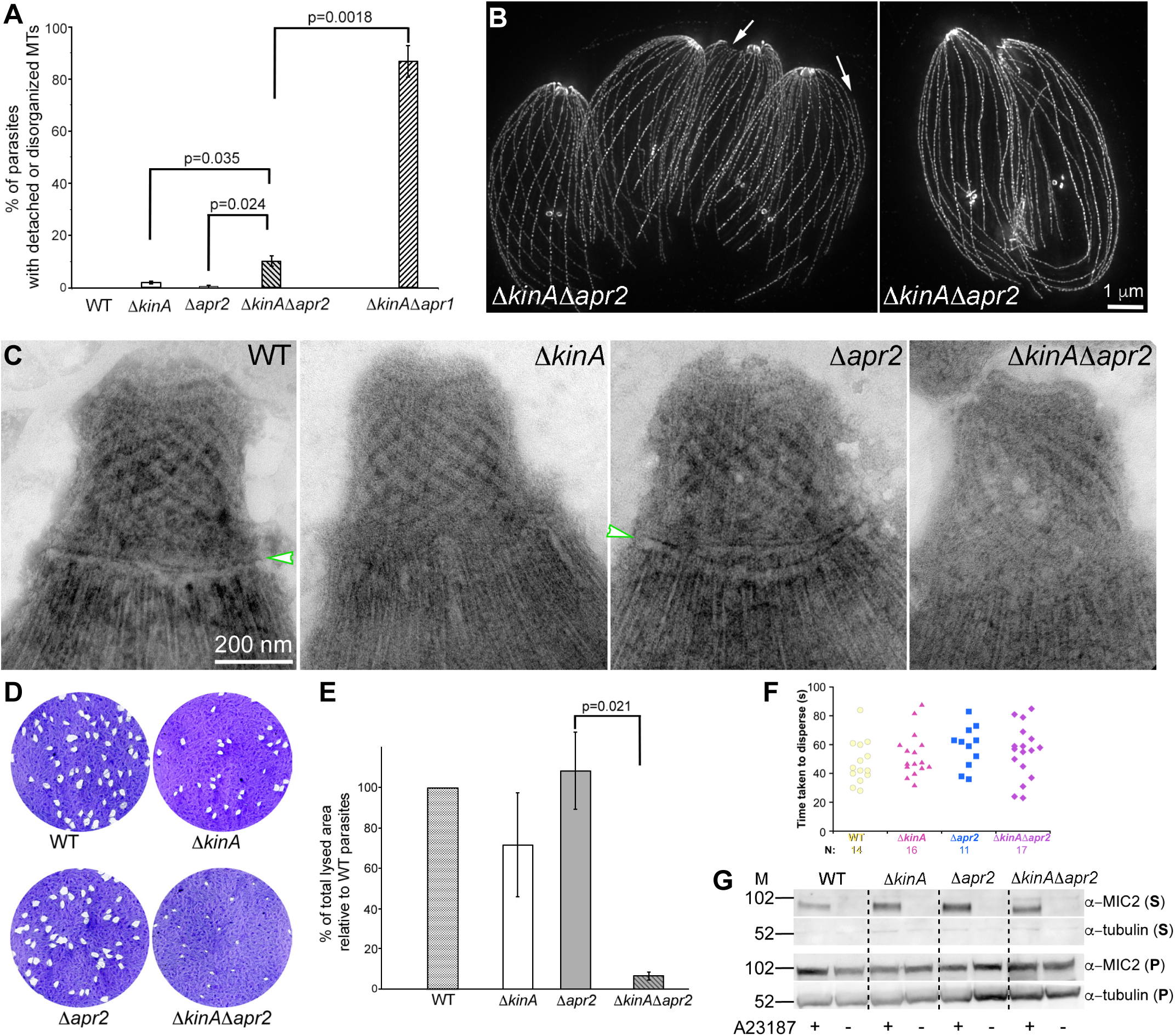
The impact of APR2 and KinesinA on the organization of the cortical MTs, the apical polar ring, and the parasite lytic cycle **A.** Bar graphs that compare % of parasites with disorganized or detached cortical MTs in the RH*Δku80* parental (WT), *ΔkinesinA* (*ΔkinA*)*, Δapr2, ΔkinesinAΔapr2,* and *ΔkinesinAΔapr1* parasites. **B.** ExM images of *ΔkinesinAΔapr2* parasites labeled with an anti-tubulin antibody as examples of normal and abnormal MT arrays. *Left* image: a vacuole that contains several parasites with detached MTs (arrows). *Right* image: twp parasites with disorganized MTs. Bar ≈ 1 µm prior to expansion. **C.** Negative-staining EM images of TX-100 extracted RH*Δku80* parental (WT), *ΔkinesinA, Δapr2,* and *ΔkinesinAΔapr2* parasites. Arrowheads indicate the annulus associated with the roots of the cortical MTs traditionally identified as the apical polar ring. This band is undetectable in *ΔkinesinA,* and *ΔkinesinAΔapr2* parasites. **D.** Plaque assays of the RH*Δku80* parental (WT), *ΔkinesinA, Δapr2*, and *ΔkinesinAΔapr2* parasites. Plaques are generated by cycles of parasite invasion, replication, and egress that destroy the host cells. **E.** Bar graphs that compare the % of the total lysed area (i.e., plaquing efficiency) relative to the WT parasites for the WT (reference), *ΔkinesinA, Δapr2,* and *ΔkinesinAΔapr2* parasites. Three independent experiments were conducted. Error bars: standard error. P-values were calculated by unpaired Student’s t-tests. **F.** Dot plots of time-taken-to-disperse after treatment with 5 µM A23187 for the RH*Δku80* parental (WT), *ΔkinesinA, Δapr2*, and *ΔkinesinAΔapr2* parasites. The datasets for the *ΔkinesinA* and *ΔkinesinAΔapr2* parasites each contained one outlier with a dispersal time higher than 140 seconds, which are not included in the dot plots. **G.** Western blots of the secreted (**S**) and pellet (**P**) fractions of RH*Δku80* parental (WT), *ΔkinesinA, Δapr2*, and *ΔkinesinAΔapr2* parasites with (+) or without (-) A23187 treatment, detected by MIC2 and tubulin antibodies. M: molecular weight markers indicated in kDa. Image contrast was enhanced and inverted to aid visualization.

To determine how the loss of KinesinA and APR2 affects the structure of the apical polar ring, we examined wild-type parental, *ΔkinesinA*, *Δapr2*, and *ΔkinesinAΔapr2* parasites by electron microscopy using negative staining after TX-100 extraction (Figure 7C). We found that the removal of APR2 alone does not have a detectable impact. In the *ΔkinesinAΔapr2* parasites, the annulus associated with the roots of the cortical MTs traditionally recognized as the apical polar ring (Figure 7C, arrowheads) is not detectable. This is likely solely due to the loss of KinesinA, because, as previously reported, the removal of KinesinA alone also results in the loss of this band.

Using plaque assays, we assessed the ability of *ΔkinesinA, Δapr2,* and *ΔkinesinAΔapr2* parasites to infect and destroy host cells relative to their parental line (Figure 7D-E). We found that the plaquing efficiency of *ΔkinesinA* and *Δapr2* is ∼ 70% and 108% of the parental RH*Δku80* (WT) line, respectively. In contrast, the plaquing efficiency of *ΔkinesinAΔapr2* is ∼ 6.6%, far lower than the ∼74% expected if the effects of the deletion of the two genes were simply additive.

Invasion and egress are two essential steps in the lytic cycle. We compared the invasion efficiency of the *ΔkinesinAΔapr2* parasite with the wild-type parental, and the single knockout parasites by using a dual-color assay that distinguishes the intra and extracellular parasite based on accessibility of the surface antigen P30 [33, 34]. We found that the invasion efficiency of *ΔkinesinAΔapr2* parasites is ∼ 34% of that of WT parasites, again indicating a synergistic effect of the knockout of KinesinA and APR2, because it is lower than the predicted invasion efficiency (0.78 × 0.66 = 51%) of the Δ*kinesinA*Δ*apr2* parasites if the effect were simply additive (Table 1). We also assessed egress efficiency of the *ΔkinesinAΔapr2* parasite by an induced-egress assay, in which the infected cultures were treated with 5 μM A23187, a calcium ionophore. We found that there is no significant difference in the speed of parasite dispersal between the *ΔkinesinAΔapr2* and the parental parasites (Figure 7F). A key process when the parasite switches between the intra and extracellular states is secretion from the membrane-bound organelles, micronemes. To determine the impact of loss of KinesinA and APR2 on microneme secretion, we examined the secretion of the major adhesin, MIC2 (Microneme Protein 2), and found that the A23187-induced MIC2 secretion of the *ΔkinesinAΔapr2* parasite remains robust (Figure 7G).

**Table 1.**
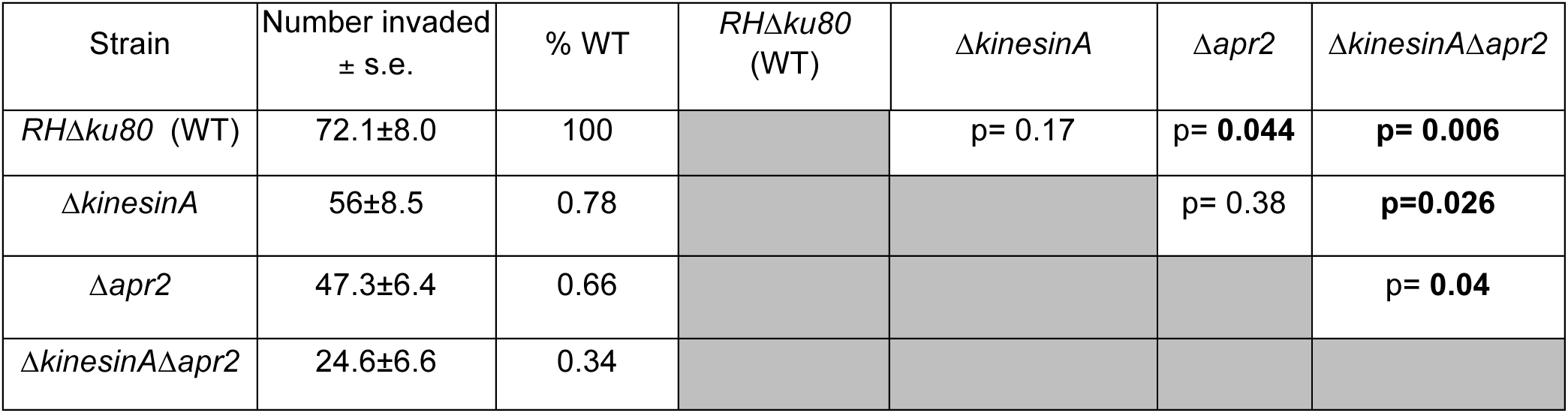
Quantification of invasion for four *T. gondii* lines. The number of intracellular parasites per field was counted in twenty fields per strain, in each of three independent biological replicates. s.e.: Standard error of the mean. P-values from unpaired Student’s t-tests are indicated on the right.

## DISCUSSION

Several components of the apical polar ring have been identified previously [22, 28, 32, 35, 36], but only KinesinA was shown to be recruited to early daughters [22]. Here we show that APR2 is another early component of the apical polar ring. The knockout of APR2 alone has no impact on the parasite lytic cycle or MT arrangement. However, it appears to act synergistically with KinesinA. The knockout of KinesinA or APR2 individually has modest or no impact, but the removal of both components results in a severely impeded lytic cycle. In a significant minority of the *ΔkinesinAΔapr2* parasites, the cortical MTs are detached from the apex. A synergistic effect was also observed when KinesinA and APR1 were both knocked out, although the MT detachment phenotype here is significantly more severe. It is worth noting that APR1 becomes detectable only in the apical polar ring of relatively late daughters [22, 27]. The knockout phenotype shows that APR1 is important for the stability of the MT-apical polar ring connection, so this late recruitment of APR1 indicates a need for additional stabilization of the MT array against the stress exerted by cortex expansion and growth as the daughter develops. Comparison of the localization of RNG1 and APR2 revealed that these proteins belong to two distinct substructures with drastically different sensitivity to perturbations. In the *ΔkinesinAΔapr1* parasite, the RNG1 labeling fragments, but the APR2 ring remains intact. The *ΔkinesinAΔapr1* parasite therefore provides a convenient genetic background for assigning various apical polar ring components to different structural domains.

As the anchoring structure for the minus ends of the cortical MTs, the apical polar ring provides a model for exploring two basic cellular processes: the formation of a ring-like cytoskeletal structure, and the patterning of a defined MT array. A well-studied example of a cytoskeletal ring-like structure is the actin-myosin based contractile ring in yeast and mammalian cells [37, 38]. This type of structure is “born” as a ring through the preferential polymerization of the cytoskeletal filaments at the division plane. Here, using APR2 as the marker, we show that the assembly of the apical polar ring proceeds incrementally. The ring precursor first appears as a short arc, which extends towards the centriole region and eventually forms a closed ring. The arc of the future apical polar ring progresses ahead of the step-wise nucleation the MT array. This pattern is consistent with the apical polar ring laying down initiation sites for subsequent MT polymerization. It was recently reported that, in addition to localizing to the centrioles as previously described [39], γ-tubulin is also located at the spindle pole and the apical polar ring region [29]. The knockdown of γ-tubulin annihilates the formation of almost all tubulin-containing structures, including the cortical MTs, the conoid, and the spindle [29]. This indicates that while the γ-tubulin complex is involved in general MT nucleation, it cannot be responsible for the differentiation of various MT-containing structures. We propose that the specificity of the origination of the cortical MT array is coded in the apical polar ring through certain scaffold proteins that recruit nucleating factors such as γ-tubulin. So far scaffold components that grossly affect the initial cortical MT patterning have not been identified. This calls for a much more thorough compilation of the composition and function of the apical polar ring.

Remarkably, the assembly of the basal complex, which is located at the opposite end of the parasite from the apical polar ring, follows essentially the same pattern both in terms of timing as well as direction. This indicates that the assembly of not only the MT array, but also the apical-basal axis is dictated by a shared responsiveness to the same signal or shared recognition of some structural gradient. Given that the assembly invariably proceeds towards the centrioles, it is conceivable that the shared element is coded in the centrioles and/or associated structures. A known structural connection between the centrioles and the daughter cytoskeletal framework is the Striated Fiber Assemblins (SFA) fiber that links the centrioles with the daughter apical complex [40]. However the impact of the fiber seems to be at the level of triggering the initiation rather than dictating the direction of the assembly, because the formation of daughter parasites is inhibited when the SFA proteins are knocked down [40].

Similar to the apical polar ring, the basal complex also grows ahead of the MT array. The basal arc fully develops into a ring before all MTs are nucleated. This suggests that the formation of the basal ring is not dependent on the development of the MT array even though they are constructed in concert. Consistent with this interpretation, we found previously that the MORN1 ring still forms when daughter MT polymerization is inhibited by oryzalin treatment [14]. Furthermore, when ectopically expressed in bacteria, MORN1 forms into rings by itself [23]. However, despite its intrinsic property to polymerize into a ring, MORN1 in *Toxoplasma* assembles only into the basal complex at the distal end of the forming daughters, where the plus-end of the cortical MTs is located. Because the ring-forming activity of MORN1 is not dependent on MT polymerization or membrane association, the specific localization of MORN1 to the basal end of the daughter is almost certainly due to bridge proteins. One would also predict that the removal of these bridge proteins should result in displaced MORN1 ring due to decoupling. So far no such proteins have been identified.

It has been proposed that the apical complex might have originated from structures associated with a flagellum [41]. The molecular link includes a number of shared components, such as centrin, dynein light chain (DLC), SAS6-like protein, and MORN-domain containing protein, as well as the SFA fiber that connects the centrosome/centrioles with the daughter apical complex [13, 24, 40, 42]. In other eukaryotes, SFA proteins form the striated rootlet that associates with the basal body. The potential evolutionary connection between the apical complex and the flagellum is further supported by the structural analysis of *Chromera*, which showed that the pseudoconoid, structural “homolog” of the conoid, is located close to the basal body of one of the two flagella [41, 43]. Interestingly, some of the proteins that suggest a molecular link between the apical complex and flagellum are also found in the basal complex, such as centrin2, DLC, and MORN1 [13, 24]. It is therefore worth considering that the basal complex is also a degenerate flagellum-related structure. If the apical and basal complexes were indeed derived from a di-flagella architecture, this could explain both their shared molecular composition as well as their highly coordinated construction.

## MATERIALS AND METHODS

### T. gondii cultures and transfection

*T. gondii* tachyzoites were cultured and transfected as described previously [22, 30, 44, 45].

### *Plasmid construction* (See Table S1 for primers used in PCR amplification)

Genomic DNA (gDNA) and Coding sequences (CDS) were prepared as described in [45]. *pTKO2_II-mEmerald-APR2 (mE-APR2)* knock-in: ∼1.9 kb fragments upstream (5’UTR) or downstream (3’UTR) of the APR2 (TGGT1_227000) genomic locus were amplified from the parasite genomic DNA by PCR using primers S1 and AS1 (5’UTR), and S2 and AS2 (for 3’ UTR) and inserted at the *Not*I (5’UTR) or *Hind*III (3’UTR) site of plasmid pTKO2-II-mCherryFP [46] using the NEBuilder HiFi Assembly kit. The CDS for mEmeraldFP-APR2 was HiFi assembled into the *AsiS*I site of this construct, generating pTKO2_II-mE-APR2. The mEmeraldFP CDS was amplified using S3 and AS3. The CDS for APR2 was amplified using primers S4 and AS4, S5 and AS5, S6 and AS6. A linker sequence coding for SGLRS was added between the APR2 and Emerald coding sequences, and the Kozak sequence from the endogenous APR2 locus (TGGTGTCAGatg) was added to the 5’ end of the mEmerald coding sequence. The backbone of the pTKO2_II-mE-APR2 plasmid contains a cassette driving expression of cytoplasmic mCherryFP, to help identify and exclude non-homologous or single homologous recombinants [46].

### Generation of knock-in, endogenously tagged, knockout, complemented, and transgenic parasites

mE-APR2 knock-in line: ∼1 x 10^7^ RHΔhxΔku80 parasites were electroporated with 40 µg of NotI linearized pTKO2_II-mE-APR2 and selected with 25 µg/mL mycophenolic acid and 50 µg/mL xanthine. Clones were screened by for mE-APR2 fluorescence and for lack of cytoplasmic mCherry fluorescence.

Clones were confirmed with genomic PCRs and then by Southern blot. Clones verified by Southern blots were used in the generation of *Δapr2* parasites.

*Δapr2* line: *mE*-*APR2* knock-in parasites were electroporated with 30 µg of pmin-Cre-eGFP_Gra-mCherry [30], selected with 80 µg/mL of 6-thioxanthine, and screened for the loss of mEmerald fluorescence. Clones were confirmed by genomic PCRs and then verified by Southern blot.

*ΔkinesinA: mE-APR2* knock-in and *ΔkinesinAΔapr1: mE-APR2* knock-in lines: *ΔkinesinA* or the *ΔkinesinAΔapr1* parasites [22] were electroporated with linearized pTKO2_II-mE-APR2 plasmid. The clones in which homologous recombination had occurred were selected and screened for as described above for the *mE-APR2* knock-in line.

*ΔkinesinAΔapr2* line: *ΔkinesinA: mE-APR2* knock-in parasites were electroporated with pmin-Cre-eGFP_Gra-mCherry. Clones were selected and screened for as described above for the *Δapr2* parasites.

RNG1-mCherry expressing *ΔkinesinAΔapr1: mE-APR2* knock-in parasites: *ΔkinesinAΔapr1: mE-APR2* knock-in parasites were electroporated with pLIC-RNG1-mCherry linearized with *Eco*RV, selected with 1 µM pyrimethamine for two passages until the population became drug resistant and cloned by limited dilution. pLIC-RNG1-mCherry was a kind gift from Dr. Naomi Morrissette at the University of California, Irvine [32].

*mE-APR2* expressing TKO parasite: *Δtlap2Δtlap3Δspm1* (”TKO”) parasites [30] were electroporated with linearized pTKO2_II-mE-APR2 knock-in plasmid and selected with 25 µg/mL mycophenolic acid and 50 µg/mL xanthine. Clones were screened for mE-APR2 fluorescence and for lack of cytoplasmic mCherry fluorescence.

### Southern blotting

The Southern blotting protocol was largely as described in [30, 46]. To probe and detect changes in the *APR2* genomic locus in the parental (RHΔ*ku80*Δ*hx*), *mE-APR2* knock-in and *Δapr2* parasites, 5 µg of gDNA from each line was digested before hybridization with a 5’UTR, CDS, or 3’UTR probe. Gel purified templates for probe synthesis were generated by restriction digestions from the pTKO2_II-mE-APR2 knock-in plasmid: *BstZ17*I and *Pme*I digestion to release the template for the 5’UTR probe; *Avr*II *and Afe*I digestion to release the template for the CDS probe; *Hind*III and *PspOM*I digestion to release the template for the 3’UTR probe.

For hybridization with the 5’UTR probe, parasite genomic DNA was digested with *Mlu*I. The predicted *Mlu*I fragment sizes are 8687 bp for the parental, 6767 bp for the knock-in, and 5146 bp for the knockout lines. For hybridization with the CDS probe, parasite genomic DNA was digested with *BstB*I. The predicted *BstB*I fragment sizes are 10994 bp and 4098 bp for the parental (*i.e.,* wild-type *apr2* locus) and 4327 bp for the knock-in. As expected, no signal was detected in the lane with *Δapr2* genomic DNA when hybridized with the CDS probe. For hybridization with the 3’UTR probe, parasite genomic DNA was digested with *Afe*I. The predicted *Afe*I fragment sizes are 11754 bp for the parental, 6617 bp for the knock-in, and 8404 bp for the knockout lines.

### Three-dimensional structured-illumination microscopy (3D-SIM)

Parasites were grown in HFF cells in glass-bottom dishes (MatTek Corporation, CAT# P35G-1.5-21-C). Before imaging, the cultures were placed in L15 imaging media [Leibovitz’s L-15 (21083-027, Gibco-Life Technologies, Grand Island, NY) supplemented with 1% (vol/vol) heat-inactivated cosmic calf serum (CCS; SH30087.3; Hyclone, Logan, UT)]. Samples were imaged with a DeltaVision OMX Flex imaging station (GE Healthcare-Applied Precision) as previously described [27]. Contrast levels were adjusted to optimize the display.

### Sample preparation and imaging for Expansion Microscopy (ExM)

ExM samples labeled with only anti-tubulin antibody were processed exactly as described in [27]. In that protocol, *Toxoplasma*-infected monolayers of HFF cells on 12mm round coverslips were pre-fixed with 3.6 % formaldehyde (made from paraformaldehyde) in PBS for 15-30 min at room temperature before fixing in the anchoring solution (2% acrylamide, 1.4 % formaldehyde, in PBS) for ExM. Under this condition, the labeling of the apical polar ring in the *mE-APR2* knockin parasites using anti-GFP antibodies was weak. This is likely due to epitope masking, because omitting the pre-fixation step or adding TX-100-extraction during the pre-fixation step both significantly improve the anti-GFP signal. The preservation of parasite morphology and the strength of the anti-GFP signal were comparable between the two methods. For the latter method, *Toxoplasma*-infected monolayers of HFF cells on 12mm round coverslips were fixed with 2.4% formaldehyde in PBS for 30 seconds followed by permeabilization with 0.4% Triton X-100 in PBS for 3 min. After washing 2 times with PBS for 5 minutes, the samples were fixed again with 2.4% formaldehyde in PBS for 20 minutes and washed 2 times for 5 minutes with PBS, followed by a second permeabilization with 0.2% Triton X100 in PBS for 10 minutes and two additional washes for 5 minutes before incubation in the anchoring solution. This method was described in [47] for the ExM protocol of labeling nuclear pore complexes.

Tubulin labeling was carried out with a mouse monoclonal anti-acetylated tubulin antibody (T6793-6-11B-1, Sigma-Aldrich, 1:250) and followed by a goat anti-mouse IgG Cy3 (115-165-166, Jackson ImmunoResearch Labs, 1:400), or goat anti-mouse IgG 488 (A-11029, Invitrogen, 1:400). Labeling of the mE-APR2 was carried out using rabbit anti-GFP antibody (Torrey Pine-TP401, 1:200), followed by goat anti-rabbit IgG Alexa488 (A-11034, Invitrogen, 1:400). Labeling of MORN1 was carried out using a rat anti-TgMORN1 antibody [23], followed by goat anti-rat IgG Cy5 (Jackson ImmunoResearch Labs 112-175-167). All other steps in ExM and antibody labeling were performed as previously described [27].

Expanded samples were imaged as described previously [27]. Sum or maximum projections were presented in the figures. Contrast levels were adjusted to optimize the display. Estimation of the expansion ratio was described previously [27].

### Electron microscopy

Suspensions of extracellular parasites were treated with A23187 and processed for negative staining as described in [45]. The negatively stained samples were imaged on a Talos TEM (Thermo Fisher) at 120 keV.

### Plaque assay

100 freshly harvested parasites per well were used to infect confluent HFF monolayers in 6-well plates. After incubation at 37°C for 7 days, the cultures were rinsed, fixed, stained, and scanned as described in [45]. Three independent experiments were performed.

### Invasion, egress, and microneme secretion assays

Immunofluorescence-based invasion assays were carried out as described in [45] with some modifications. ∼1 x 10^6^ freshly egressed parasites were used to inoculate a MatTek dish of nearly confluent HFF cells. After 10 min incubation on ice and then 60 min incubation at 37°C, the samples were processed for immunofluorescence as described in [45]. Three biological replicates were performed. Parasites in 20 fields were counted for each strain per biological replicate. In total (intracellular + extracellular), 6230 parasites were counted for the RH*Δhx* parent, 4303 for *ΔkinesinA*, 4616 for *Δapr2*, 2877 for *ΔkinesinAΔapr2* parasites. P-values were calculated by unpaired Student’s t-tests.

The induced egress assays were carried out as described in [45]. Microneme secretion assays were carried out as described in [31].

## Acknowledgments

We thank Matea Susac for tissue culture support, and Isadonna F. Tengganu for helpful discussions.

---

**Table S1.**
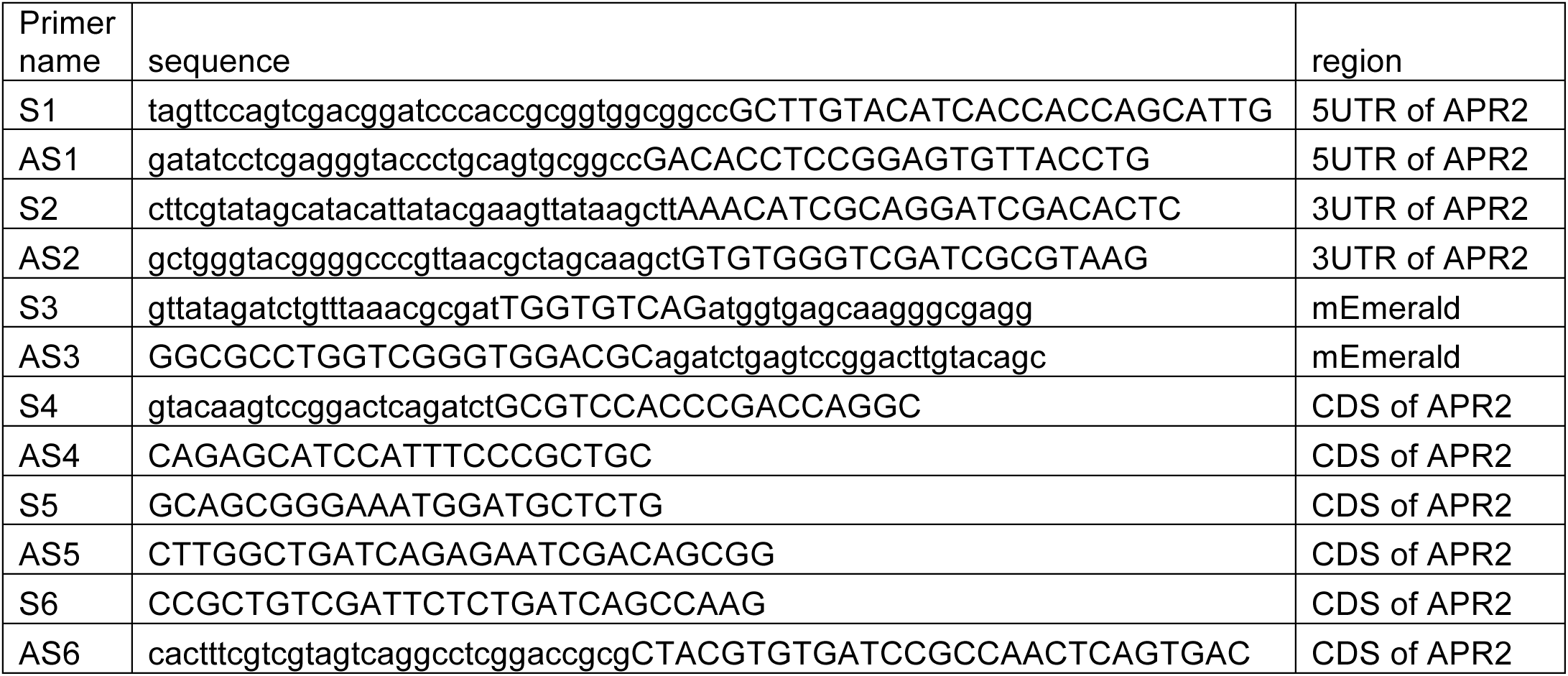

